# Exploring the Dynamics of Follicle Development and Hormone Synthesis: The Role of Oxygen Tension in Rhesus Macaque Follicle Culture

**DOI:** 10.1101/2025.05.06.652505

**Authors:** Kang Wang, Shally Wolf, Mary B. Zelinski, Adam J. Krieg

## Abstract

In vitro culture of cryopreserved ovarian follicles has the potential to extend fertility options for young women seeking to preserve ovarian tissue prior to undergoing cancer treatments. Successful implementation of this strategy has been elusive and likely requires a more complete understanding of the microenvironment of the developing ovarian follicles, including ovarian oxygen concentrations. The oxygen tension within the reproductive tract plays a crucial role in follicular development and oocyte maturation. While *in vitro* culture systems often use atmospheric oxygen (21%), the native environment *in vivo* is significantly lower, ranging from 1.5% to 8.7%. This study aimed to investigate the effects of reduced oxygen tensions (3% and 5%) on follicle survival, growth, antrum formation, and hormone production in cultured secondary follicles from rhesus macaques (*macaca mulata*). A total of 300 follicles were isolated from 7 animals and cultured under three oxygen conditions: 3%, 5%, and 21% O_2_. Follicle survival and antrum formation were assessed weekly by microscopy and Kaplan–Meier survival analysis, while growth dynamics and hormone levels (estradiol, progesterone, AMH, and inhibin B) were monitored throughout the culture period.

Results demonstrated that follicles cultured at 3% and 5% oxygen exhibited significantly higher survival rates and antrum formation compared to those cultured at 21% O_2_. No significant differences in survival were observed between the 3% and 5% oxygen groups. Growth dynamics revealed distinct patterns, with both low oxygen groups promoting more robust and sustained follicle growth, while atmospheric oxygen led to rapid degeneration. Hormonal analysis showed that follicles in 21% O_2_ had elevated early hormone production but exhibited reduced long–term viability. In contrast, 3% and 5% oxygen delayed hormone production, reflecting a more stable and sustained follicular environment.

These findings underscore the importance of low oxygen tensions in mimicking the physiological conditions of the reproductive tract, improving follicular development, and supporting optimal hormonal function *in vitro*. This study suggests that further reducing oxygen levels to 3% may offer additional advantages for long–term follicle viability and function in reproductive technologies.

**Graphical Abstract:** 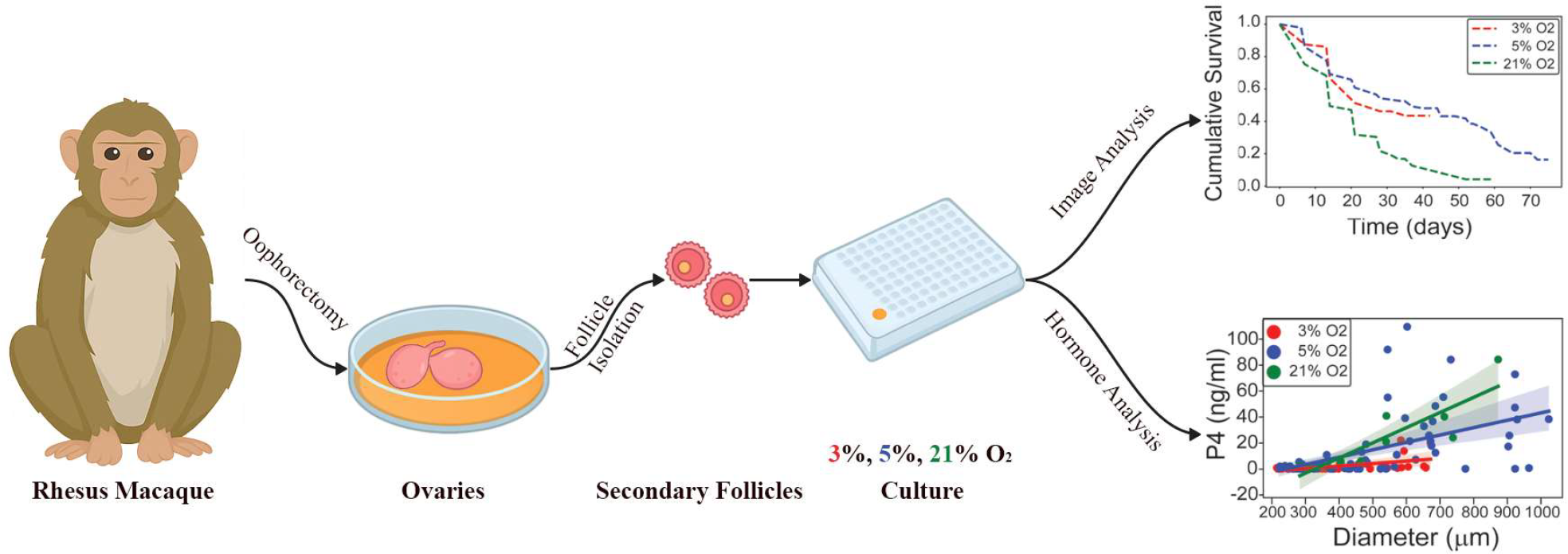

## INTRODUCTION

Recent advances in the treatment of pediatric cancers have resulted in tremendous increases in patient survival, with approximately 85% of pediatric cases surviving more than 5 years after diagnosis [1]. However, this unprecedented accomplishment also introduces unforeseen consequences: The downstream effects of chemotherapeutic agents increase the risks of chronic organ damage and dysfunction, especially for reproductive organs [2]. Because of the central role of the ovary in establishing and maintaining reproductive potential, cardiac health, and bone health in adult women, there is a great deal of interest in developing more effective methods for preserving ovarian tissues in order to restore or extend ovarian function in both pediatric cancer patients and women of advancing reproductive age [2]. Cryo–preservation of ovarian tissues followed by *in vitro* culture of secondary follicles represents a promising strategy to provide pediatric cancer patients with an *ex vivo* reserve of viable oocytes for *in vitro* culture and eventual fertilization. However, in order to ensure that *in vitro* ovarian culture techniques eventually yield viable oocytes suitable for fertilization, it is essential to optimize culture techniques to maximize *in vitro* survival and developmental potential.

Mammalian oocytes were first matured *in vitro* in the 1930s, representing a pivotal advancement in reproductive biology by enabling controlled investigating into conditions optimal for oocyte maturation [3]. Over the years, studies have demonstrated that oxygen plays a crucial role in this process, particularly in nuclear maturation – the process by which oocytes resume meiosis and reach developmental competence – which is highly sensitive to oxygen tension [4]. Early studies identified detrimental effects of atmospheric oxygen levels (21% O_2_) on oocyte development, prompting further investigation into lower oxygen tensions. These efforts emphasized the need to replicate *in vivo* conditions, characterized by significantly lower oxygen concentration, to enhance *in vitro* culture outcomes.

*In vivo* studies have shown that oocytes experience significantly lower oxygen tensions in the reproductive tract, ranging from approximately 1.5% and 8.7% O_2_, depending on species and tissue type [5-7]. In rhesus monkeys, for example, oxygen concentrations in the uterine lumen remain consistently around 1.5% throughout the menstrual cycle [5]. This contrasts sharply with the 21% atmospheric oxygen typically used in laboratory settings, a condition known to impair oocyte development due to increased oxidative stress and altered cellular metabolism [8].

Building on this foundation, culture systems utilizing 5% oxygen have consistently demonstrated improved outcomes for follicle development and oocyte maturation compared to atmospheric oxygen conditions. Studies in rhesus macaques, mice and sheep have shown that culturing preantral follicles under 5% oxygen not only supports higher survival rates but also leads to improved oocyte quality and developmental outcomes when compared to higher oxygen tensions [9-12]. Additionally, 5% oxygen has been shown to significantly enhance the maturation competence of mouse oocytes, supporting their developmental potential, while higher oxygen concentrations have been found to impair these processes [8]. Similar benefits have been observed in buffalo and porcine species, where culturing oocytes at 5% oxygen not only increased maturation efficiency but also enhanced the developmental competence of early embryos [13, 14]. These findings underscore the physiological relevance of low oxygen conditions, emphasizing the importance of replicating the *in vivo* environment to optimize *in vitro* follicle development and oocyte maturation.

Lower oxygen tensions further support oocyte maturation by reducing oxidative stress, thereby directly improving follicle survival, oocyte quality, and developmental competence. For example, culturing follicles at 5% oxygen has been linked to reduced rates of oxidative stress and apoptosis in granulosa cells, thereby improving follicle survival and function [15]. In rhesus monkeys, embedded secondary follicle culture at 5% O_2_ improved survival rates compared to atmospheric oxygen conditions [9].

However, while 5% O_2_ has emerged as the preferred condition for many culture systems, the native oxygen environment in the reproductive tract may be even lower, raising the question of whether further reductions in oxygen tension could offer additional advantages by minimizing oxidative stress and promoting more physiologically relevant culture conditions. Culturing granulosa cells in 1% O_2_ supports the viability and function of granulosa cells *in vitro*, inducing changes in cellular phenotype and function without significantly affecting cell viability [16, 17]. Murine studies of follicle activation have also implicated reduced oxygen and HIF signaling in maintenance of quiescence [18], while analysis of human granulosa and ovarian tissues has shown that hypoxia–inducible histone demethylases are also robustly expressed in smaller follicles [19]. Collectively, these data suggest that follicles and their supporting cells may be adapted to very low oxygen tensions. In this study, we sought to explore the impact of culturing follicles at 3% oxygen, hypothesizing that this lower oxygen tension might confer additional advantages for follicle survival, growth, and oocyte maturation. To test this, we compared follicle cultures at three different oxygen tensions – 3%, 5%, and 21% – and assessed follicle survival, antrum formation, and hormone production across these conditions.

## MATERIALS AND METHODS

### Animals and ovary collection

The general care and housing of rhesus macaque monkeys were provided by the Division of Animal Resources at the Oregon National Primate Research Center (ONPRC). Animals were pair–caged in a temperature–controlled (22 °C), light–regulated rooms with a 12–hour light/dark cycle (12 L:12 D). Diet consisted of Purina monkey chow (Ralston–Purina, Richmond, IN, USA) provided twice daily, supplemented once daily with fresh fruit or vegetables, and water *ad libitum*. Animal care followed the National Institutes of Health Guide for the Care and Use of Laboratory Animals, and all protocols were approved by the ONPRC Institutional Animal Care and Use Committee.

Ovaries were collected via surgical ovariectomy from adult female rhesus macaques (n=7; aged 5–12 years). Immediately following removal, ovaries were placed into Hepes–buffered holding media (CooperSurgical, Inc., Trumbull, CT, USA) supplemented with 0.2% (v/v) human serum protein supplement (SPS, CooperSurgical, Inc.) and 10 µg/ml gentamicin (Sigma–Aldrich, St Louis, MO, USA)

### Follicle isolation and culture

Follicles were dissected according to previously described methods by Xu et al. [9, 20, 21]. Briefly, ovarian cortex was sectioned into ∼500 µm-thick slices using a Stadie-Riggs tissue slicer (Thomas Scientific, Swedesboro, NJ, USA). These cortical slices were further cut into ∼0.5 × 0.5 × 0.5 mm cubes and incubated in 6 ml holding media at 37 °C. Follicles were mechanically isolated in holding media using 31–gauge needles. Secondary follicles measuring 150 – 250 µm in diameter were selected based on: (i) intact basement membrane, (ii) two to three layers of granulosa cells, and (iii) a healthy, round, centrally located oocyte without vacuoles or dark cytoplasm. Follicles (n= 300 from 7 monkeys) were divided, and each follicle was transferred to individual wells of 96–well U-bottom, ultra-low attachment plates (Costar REF7007, Corning Incorporated, Kennebunk, ME, USA) containing 200 µl alpha minimum essential medium (αMEM, Invitrogen) supplemented with 6% (v/v) SPS, 5 µg/ml insulin, 5 µg/ml transferrin and 5 ng/ml sodium selenite (Sigma–Aldrich). While we aimed to distribute follicles from each animal across all three oxygen conditions (3%, 5%, and 21% O_2_), the number of available secondary follicles per animal varied. In cases where follicle yield was limited, some animals contributed follicles to only one or two oxygen groups. Cultures were maintained at 37 °C and 5% carbon dioxide under three oxygen tensions: 3% (v/v) O_2_ in a hypoxic glove–box chamber (Coy Laboratory Products, Grass Lake, MI, USA), 5% (v/v) O_2_ in a tri-gas incubator (Thermo), or a 20% (v/v) O_2_ in a tri-gas incubator (Thermo). Due to concerns regarding maintenance of a sterile environment during prolonged incubations, 3% O_2_ cultures were conducted for 7 weeks, whereas 5% and 21% O_2_ cultures extended longer as needed to reach experimental endpoints. For hypoxic cultures, oxygen was displaced with medical–grade nitrogen gas. Half of the culture media (100 µl) was replaced every other day, with collected media stored at –20 °C for hormonal analysis.

### Follicle survival and growth

Follicle survival, diameter and antrum formation at 5% and 21% O_2_ were assessed weekly using an Olympus CK40 inverted microscope and an Olympus DPI I digital camera (Olympus Imaging America Inc., Center Valley, PA, USA). To prevent reoxygenation of follicles cultured at 3% O_2_, images were captured within the hypoxia chamber using a Lumiscope 620 microscope (Etaluma, San Diego, CA, USA). Images were analyzed in ImageJ 1.42 (National Institutes of Health, Bethesda, MD, USA). Follicle diameter was determined from two perpendicular measurements (widest and orthogonal), averaged to provide the reported diameter. Follicles were classified as dead if the oocyte was extruded or showed clear signs of atresia, defined as darkened or collapsed oocytes no longer fully surrounded by granulosa cells.

### Ovarian steroids and AMH assays

Hormones in culture media were analyzed by the Endocrine Technology Support Core at ONPRC. Estradiol (E2) and progesterone (P4) were measured using an Immulite 2000 chemiluminescence–based automatic clinical platform (Siemens Healthcare Diagnostics, Deerfield, IL, USA). AMH and inhibin B concentration were measured by ELISA (Ansh Labs and Diagnostic Systems Laboratory, Webster, TX, USA, respectively) according to manufacturers’ protocols.

### Statistical analyses

Statistical analyses were performed in Python (python.org) using the statsmodels library [22]. Data are presented as mean ± SEM, and the ensembled averages were compared statistically using one–way ANOVA followed by the Tukey’s HSD *post hoc* test or Mann–Whitney U test for single time points. Log–rank tests were performed for Kaplan–Meier based follicle survival and antrum formation analysis. To evaluate the relationship between follicle diameter and hormone secretion (e.g. E2, P4, AMH, Inhibin B), ordinary least squares (OLS) linear regression was applied separately within each oxygen condition (3%, 5%, and 21%). Only fast-grow follicles were included in these analyses to focus on follicles that successfully activated and progressed through development. Since follicle diameter increases monotonically over time and serves as a proxy for developmental progression, and because the goal was to examine population-level trends rather than individual follicle trajectories, OLS regression was deemed appropriate despite repeated measures within follicles. Where appropriate, polynomial regression models (up to fourth degree) were also fitted and compared to linear models using adjusted R^2^, individual coefficient *p*-values, and ANOVA F-tests. For AMH, Gaussian Mixture Modeling (GMM) was used to assess whether the distribution of AMH secretion levels could be better explained by two latent subpopulations [23]. GMM models with one and two components were compared using the Bayesian Information Criterion (BIC), with a ΔBIC > 10 interpreted as strong evidence supporting bimodality [24]. P < 0.05 was accepted as statistically significant.

## RESULTS

### Culturing follicles in reduced oxygen tensions improved survival *in vitro*

In this study, we first assessed the impact of oxygen tension on follicle survival using Kaplan–Meier survival analysis over approximately 10 weeks. A total of 300 secondary follicles isolated from 7 rhesus macaques were allocated to three oxygen conditions: 3%, 5%, and 21% O_2_. The resulting survival curves revealed distinct survival trajectories for each oxygen tension level, where each downward step represents follicle degeneration events (Figure 1).

**Figure 1.**
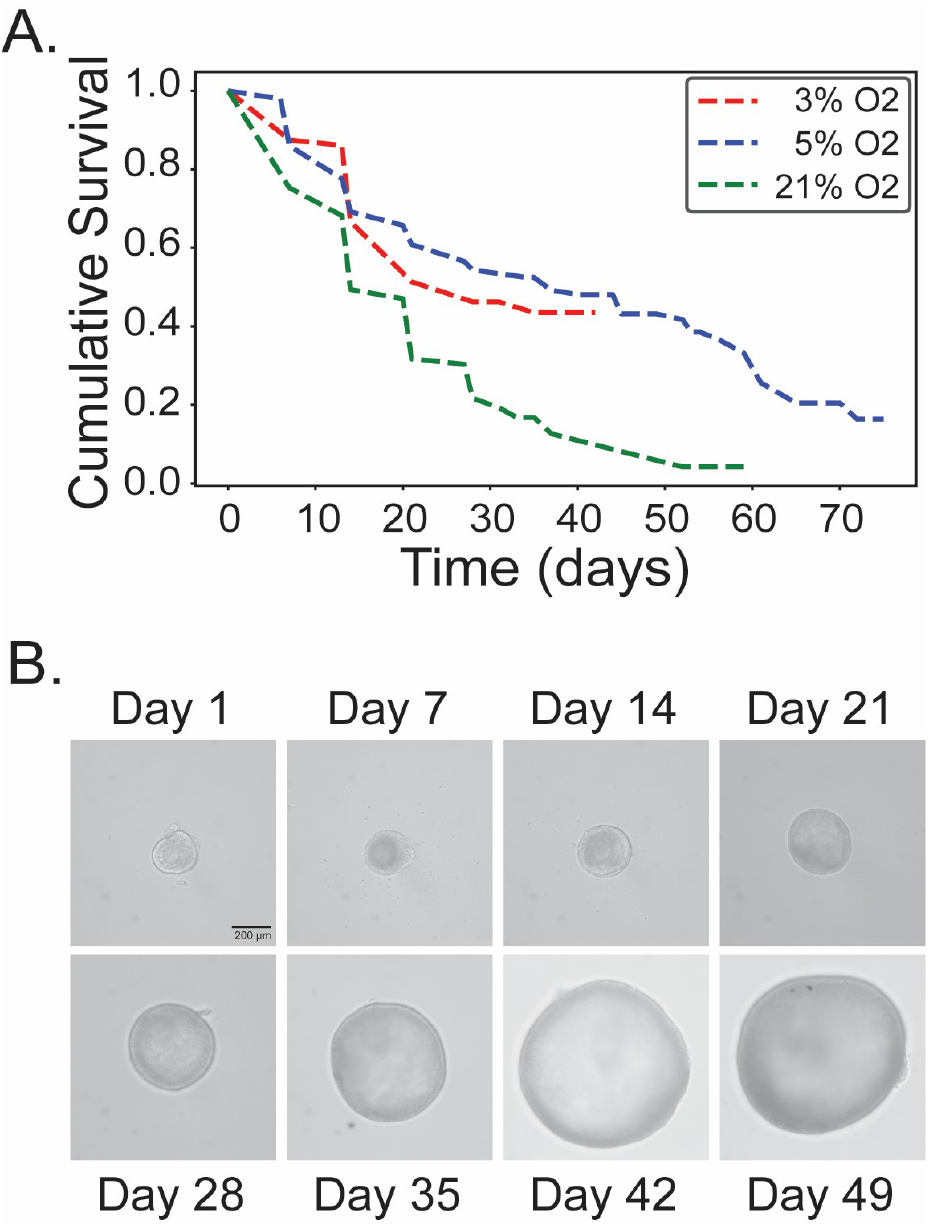
Kaplan–Meier survival curves for follicles cultured under various oxygen tensions. (**A**) The red, blue, and green dashed lines represent follicles cultured in 3% O_2_, 5% O_2_, and 21% O_2_ respectively, with each step down indicating a follicle degeneration event over a period of approximately 10 weeks. Log–rank test results indicate significant differences in survival between the 3% O_2_ and 21% O_2_ groups (*** p < 0.001), and the 5% O_2_ and 21% O_2_ groups (*** p < 0.001), while there is no statistically significant difference between the 3% O_2_ and 5% O_2_ groups (p = 0.467). (**B**) Representative images of a follicle cultured in 5% O_2_ at multiple time points demonstrating follicle survival. Scale bar (black) = 200 μm.

In the low oxygen tension group (3% O_2_), follicles displayed a gradual decline in survival, with the most significant drop observed between days 10 and 20, followed by a steady rate of degeneration. The culture in 3% O_2_ was stopped at week 7, as explained in the material and methods section. The intermediate oxygen tension group (5% O_2_) exhibited a similar initial decline but showed a slight increase in the survival rate post–day 20, maintaining a more constant survival rate until approximately day 50, where a more pronounced decline was observed. In contrast, follicles cultured at atmospheric oxygen tension (21% O_2_) showed markedly inferior outcomes, characterized by a sharp initial decline within the first 10 days, followed by continued rapid degeneration until day 20. Afterward, the degeneration rate decreased slightly, but overall survival remained significantly lower than observed in 3% or 5% O_2_.

Log–rank analysis revealed statistically significant differences in survival between the 3% O_2_ and 21% O_2_ groups (p < 0.001), as well as between the 5% O_2_ and 21% O_2_ groups (p < 0.001). Conversely, no significant difference was found between the 3% and 5% O_2_ conditions (p = 0.467), indicating that reduced oxygen tensions may similarly enhance follicle viability. Taken together, these results suggest that oxygen tension is a critical factor in follicle survival *in vitro*, with lower oxygen tensions (3% O_2_ and 5% O_2_) being more conducive to follicle longevity than atmospheric oxygen levels.

### Culturing follicles in reduced oxygen tensions increases the formation of antral follicles *in vitro*

Reduced oxygen tensions also significantly improved the proportion of follicles that underwent antrum formation (Fig. 2A). Follicles cultured at lower oxygen tensions (3% and 5% O_2_) formed antra at markedly higher proportions (45.8% and 46.9%, respectively) compared to follicles cultured under atmospheric conditions (21% O_2_), where only 17.6% formed antra. Chi-square analysis revealed statistically significant differences in antrum formation when comparing follicles cultured at 3% O_2_ vs. 21% O_2_ (p < 0.001), and 5% O_2_ vs. 21% O_2_ (p < 0.001). No significant difference was detected between the 3% and 5% O_2_ groups (p = 1.0).

**Figure 2.**
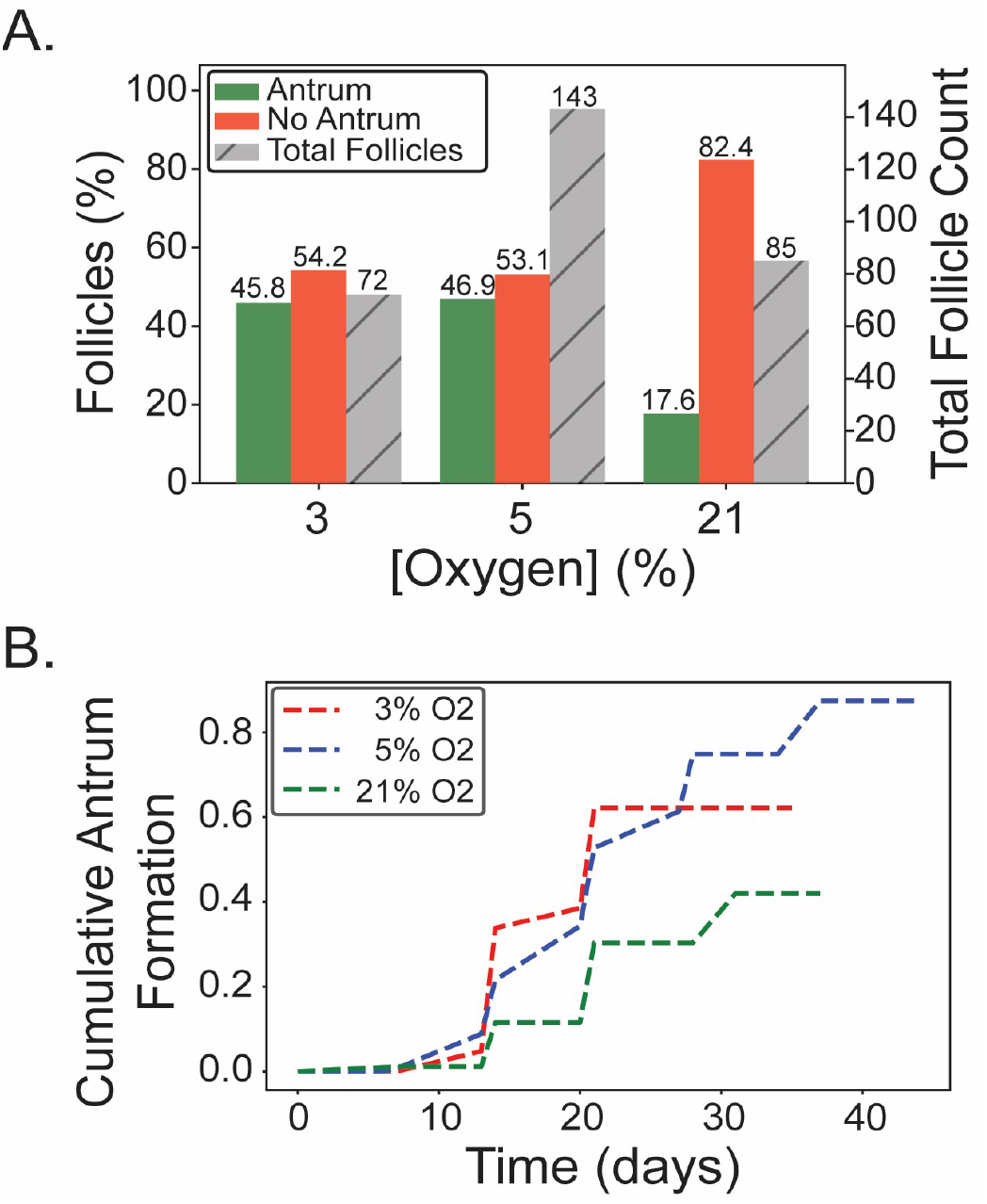
The effects of oxygen tension on follicle antrum formation, represented as percentages of the total follicle count (*A*) and the Kaplan–Meier curves for follicles antrum formation (*B*). (**A**) The primary y–axis (left) shows the percentage of follicles with antrum formation (green) or without antrum formation (red) across different oxygen tensions (3%, 5%, and 21%). The secondary y–axis (right) represents the total number of follicles, indicated by hatched bars. Chi–square test results indicate significant differences in follicle antrum formation distribution between the 3% O_2_ and 21% O_2_ groups (*** p < 0.001), and the 5% O_2_ and 21% O_2_ groups (*** p < 0.001), while there is no statistically significant difference between the 3% O_2_ and 5% O_2_ groups (p = 1.0). (**B**) Kaplan–Meier curves of antrum formation for follicles cultured under various oxygen tensions. The red, blue, and green dashed lines represent follicles cultured in 3% O_2_, 5% O_2_, and 21% O_2_ respectively, with each step up indicating a follicle antrum formation event over a period of approximately 6 weeks. Log–rank test results indicate significant differences in follicle antrum formation between the 3% O_2_ and 21% O_2_ groups (*** p = 0.001), and the 5% O_2_ and 21% O_2_ groups (*** p < 0.001), while there is no statistically significant difference between the 3% O_2_ and 5% O_2_ groups (p = 0.529).

Kaplan–Meier analysis further supported these observations by illustrating cumulative probabilities of antrum formation over time (Fig. 2B). Both low-oxygen groups (3% and 5% O_2_) demonstrated a steady increase in antrum formation rates, primarily occurring between culture days 20 and 30. In contrast, the 21% O_2_ group exhibited delayed onset and a substantially lower cumulative rate of antrum formation. Log-rank analysis confirmed significant differences in the timing of antrum formation between the 3% O_2_ and 21% O_2_ conditions (p = 0.001), as well as between the 5% O_2_ and 21% O_2_ conditions (p < 0.001). However, the timing of antrum formation did not significantly differ between the 3% and 5% oxygen conditions (p = 0.529).

These results clearly demonstrate that culturing follicles at lower oxygen tensions significantly promotes antrum formation, whereas atmospheric oxygen conditions adversely affect this important developmental milestone. Thus, oxygen tension represents a critical factor influencing follicle development and maturation in vitro.

### Impact of oxygen tension on follicle growth dynamics

To further elucidate the relationship between oxygen tension and follicular development, we characterized growth patterns of the 194 follicles that survived throughout the culture period (Fig. 3). Follicle growth trajectories were classified into categories previously described [21]: No–grow (NG, no increase in size after 28 days), Slow–grow (SG, diameter increase less than 2.5-fold after 28 days), and Fast–grow (FG, diameter increase greater than 2.5-fold after 28 days. In addition, we identified a previously undescribed category unique to follicles cultured at 5% O_2_, termed “delayed Fast-grow” (dFG). These follicles initially displayed SG growth kinetics but transitioned to FG kinetics after approximately 6 weeks of culture.

**Figure 3.**
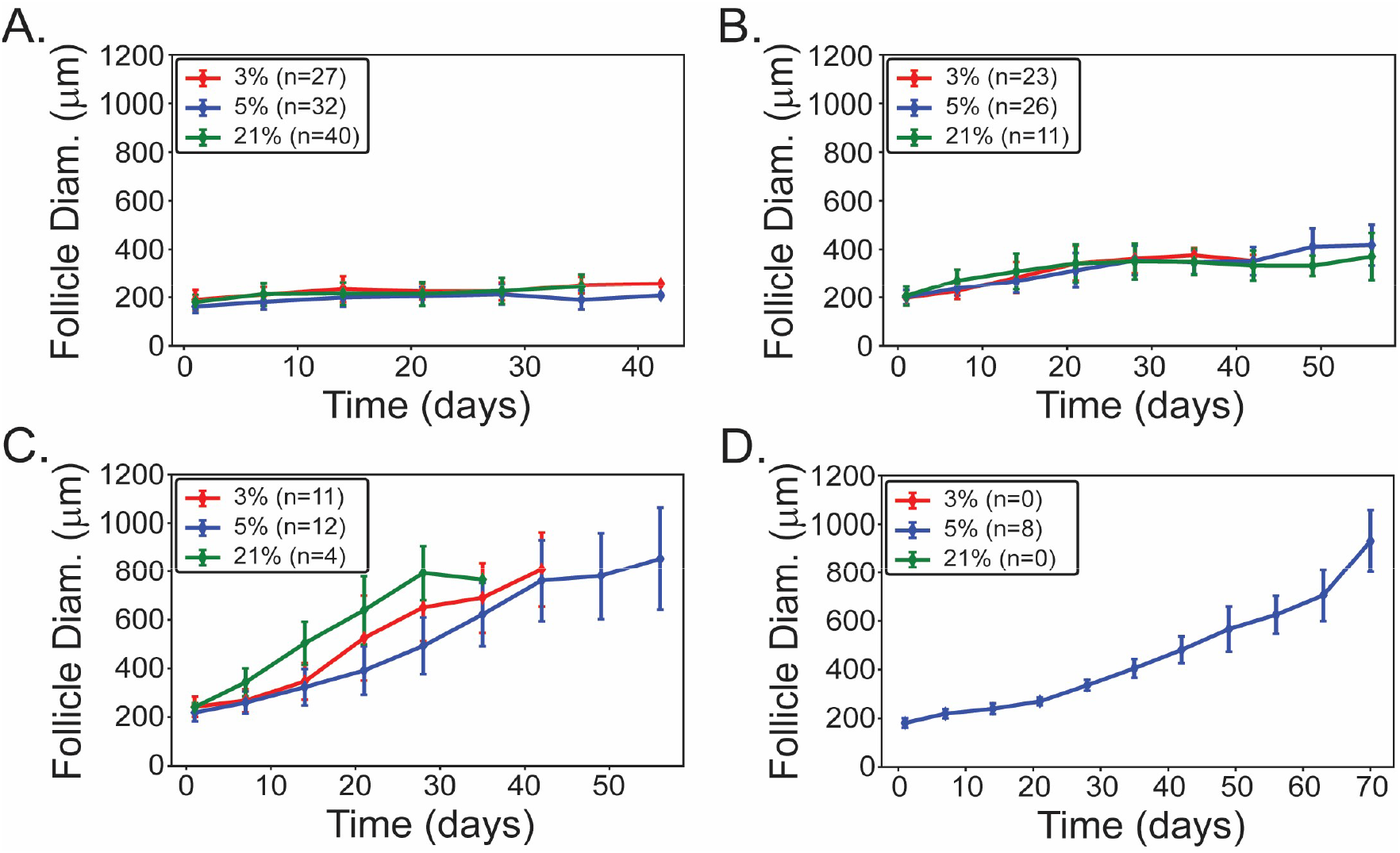
Follicle growth trajectories under difference oxygen tensions. Follicle growth patterns were classified as (**A**) No-grow (NG; no increase in diameter after 28 days), (**B**) Slow-grow (SG; less than 2.5-fold increase after 28 days), (**C**) Fast-grow (FG; greater than 2.5-fold increase after 28 days), and (**D**) delayed Fast-grow (dFG; initially SG kinetics shifting to FG after ∼6 weeks). Mean follicle diameters (μm) are presented over the culture duration for follicles cultured at 3% (red), 5% (blue), and 21% O_2_ (green). Error bars represent the standard error of the mean (SEM), with sample sizes (n) indicated.

In contrast to significant oxygen-dependent effects observed in survival and antrum formation, oxygen tension did not significantly alter follicle diameter within each individual growth category. However, the proportion of follicles assigned to each growth category varied significantly according to oxygen level (Fig. 4). Specifically, the proportion of follicles categorized as Fast–grow (FG) and delayed Fast–grow (dFG) was significantly higher under 3% and 5% O_2_ compared to 21% O_2_, whereas a larger fraction of No–grow (NG) and Slow–grow (SG) follicles was observed under atmospheric oxygen conditions (21% O_2_; p = 0.008 and p = 0.001, respectively).No statistically significant difference was observed between the 3% O_2_ and 5% O_2_ groups (p = 0.083). While differences in the maximum follicle diameter achieved were not statistically significant, follicles cultured under hypoxic conditions tended to achieve their peak diameters over extended culture periods compared to those cultured in atmospheric conditions (Fig. 4B–D).

**Figure 4.**
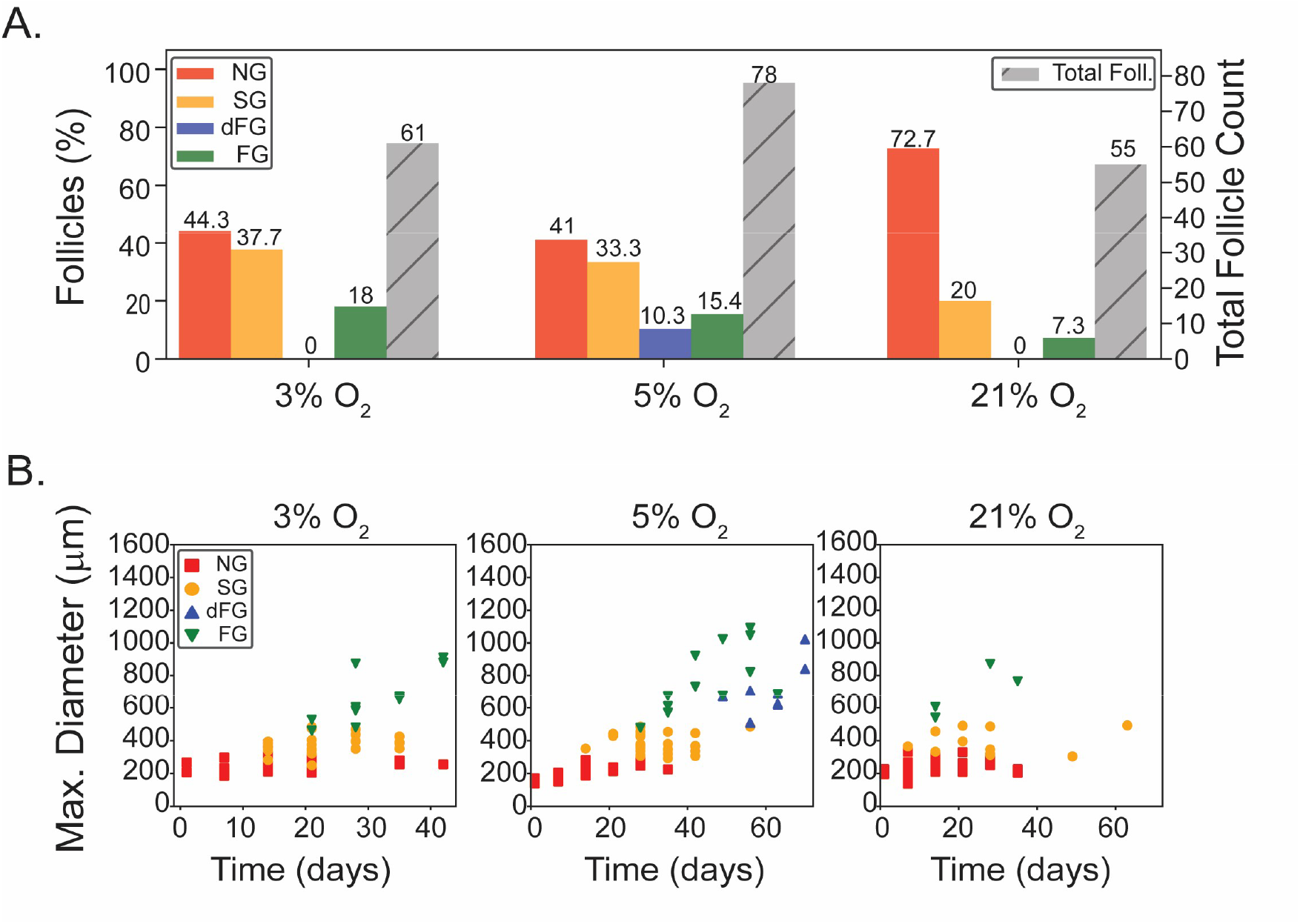
Distribution of follicle growth pattern by oxygen tension groups, represented as percentages of the total follicle count (*A*) and the max follicle growth diameter (*B, C, D*). (**A**) The primary y–axis (left) shows the percentage of follicles in each category: No–grow (NG, red), Slow–grow (SG, yellow), delayed Fast– grow (dFG, blue), and Fast–grow (FG, green) across different oxygen tensions (3%, 5%, and 21%). The secondary y–axis (right) represents the total number of survival follicles, indicated by hatched bars. Chi– square test results indicate significant differences in follicle categorical distribution between the 3% O_2_ and 21% O_2_ groups (* p = 0.008), and the 5% O_2_ and 21% O_2_ groups (*** p = 0.001), while there is no statistically significant difference between the 3% O_2_ and 5% O_2_ groups (p = 0.083). (**B–D**) Scatter plot of maximum growth size of individual follicle during culture under different oxygen tensions. The y–axis represents the maximum diameter of follicles in micrometers, and the x–axis represents time in days when each follicle reaches its maximum size. The plots are separated into three panels corresponding to oxygen levels of 3%, 5%, and 21% from left to right.

These data indicate that lower oxygen tensions (3% and 5% O_2_) support improved follicle growth potential, manifested by more favorable categorical distributions and sustained growth trajectories compared to atmospheric oxygen. These findings underscore oxygen tension as a critical determinant in optimizing follicle culture environments to enhance follicular growth and developmental outcomes.

### Hypoxic culture conditions change the magnitude and dynamics of hormone production *in vitro*

We next evaluated the influence of oxygen tension on the production dynamics of key reproductive hormones, including Estradiol (E2), Progesterone (P4), Anti-Müllerian Hormone (AMH), and Inhibin B (INHB), across different oxygen conditions (Fig. 5). Hormone concentrations were analyzed separately for Slow-grow (SG) and Fast-grow (FG) follicles.

**Figure 5.**
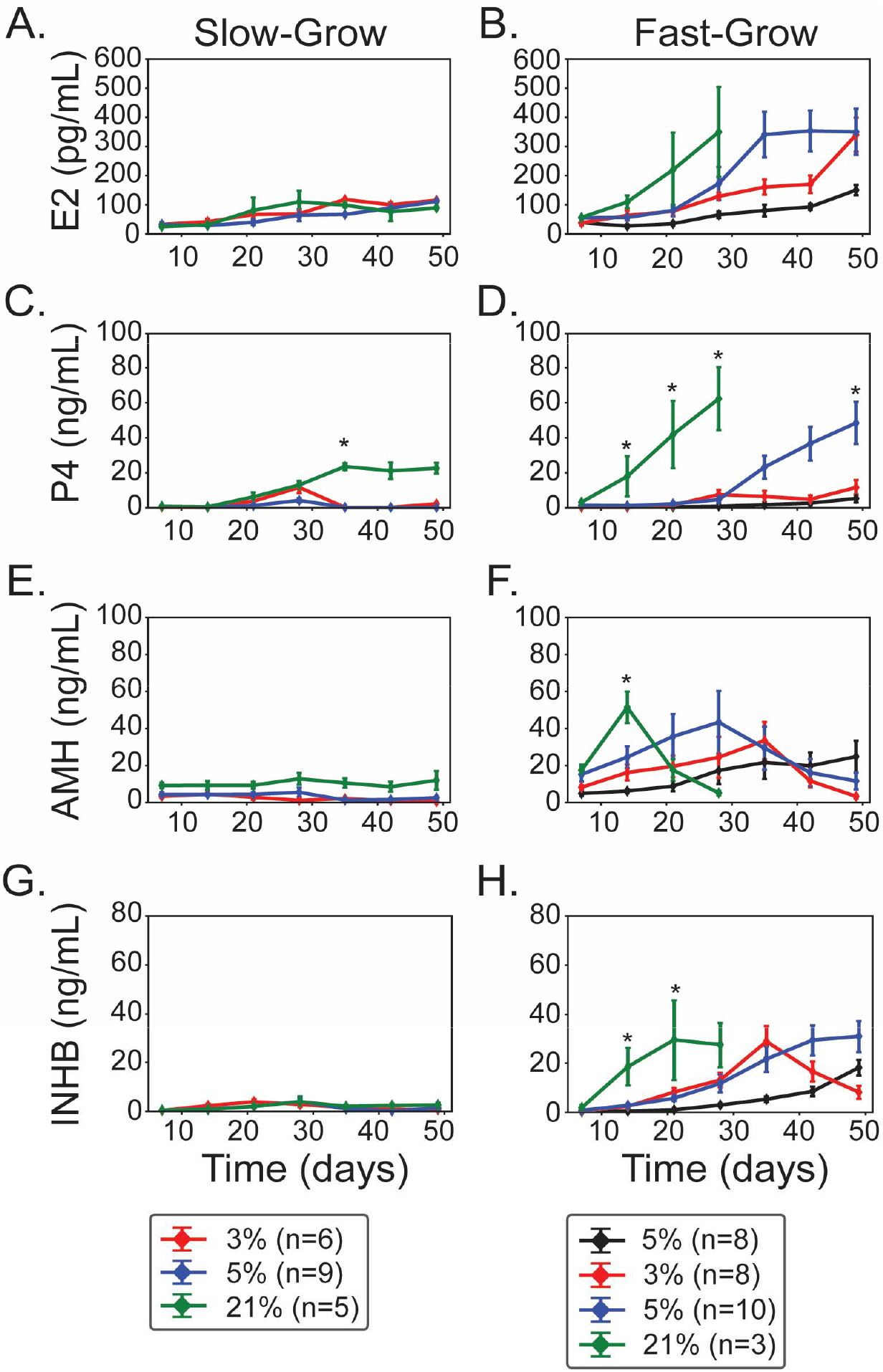
Temporal patterns of hormone production by follicles cultured under different oxygen tensions. Panels show mean hormone concentrations over seven weeks for Estradiol (**A–B**), Progesterone (**C–D**), AMH (**E–F**), and Inhibin B (**G–H**) separately for Slow-grow (SG; left column) and Fast-grow (FG; right column) follicles cultured under 3% O_2_ (red), 5% O_2_ (blue), and 21% O_2_ (green). Black lines in fast–grow panels indicate delayed growth (dFG) under 5% O_2_. Asterisks denote significant differences between culture conditions (p < 0.05). ‘n’ represents the number of follicles; error bars show SEM.

In SG follicles, mean E2 levels remained relatively stable across all oxygen tensions, without statistically significant differences (ANOVA: all time points, p > 0.05; Fig. 5A). Conversely, in FG follicles E2 levels at 21% O_2_ increased significantly early in culture (within the first 4 weeks), correlating with their rapid initial growth (ANOVA: day 14: p = 0.028; day 21: p = 0.0008; day 28: p = 2.6e-05; Fig. 5B). Notably, due to the limited survival of FG follicles at 21% O_2_ beyond four weeks, media samples from this group were only collected up to day 28. FG follicles cultured at lower oxygen levels (3% and 5% O_2_) displayed a delayed yet sustained increase in E2 production compared to follicles cultured at 21% O_2_.

Progesterone production in SG follicles was significantly elevated at later time points under 21% O_2_ (day 21: p = 0.0002; day 28: p < 0.0001), indicating enhanced steroidogenic activity under atmospheric conditions (Fig. 5C). Similarly, FG follicles at 21% O_2_ demonstrated rapid, pronounced increases in progesterone within the initial 4 weeks of culture (day 14: p = 0.028; day 21: p = 0.0008), correlating strongly with their accelerated growth dynamics (Fig. 5D). In contrast, FG follicles cultured at 5% O_2_ significantly elevated progesterone levels only after seven weeks of culture, whereas those cultured at 3% O_2_, as well as dFG follicles at 5% O_2_, showed minimal or no progesterone elevation (Fig. 5D).

AMH concentrations in SG follicles were relatively stable across all conditions but significantly elevated at 21% O_2_ early in culture (day 7: p = 0.097; day 14: p = 0.297), suggesting minor yet discernible sensitivity to atmospheric oxygen (Fig. 5E). FG follicles cultured at 21% O_2_ showed a sharp AMH peak at week 2, aligning with previous observations [9] whereas those cultured under hypoxic conditions (3% and 5% O_2_) had delayed and broader AMH peaks without significant differences at later time points (ANOVA: all later points, p > 0.05; Fig. 5F).

Inhibin B levels showed minimal variation in SG follicles across all oxygen tensions (ANOVA: all later points, p > 0.05; Fig. 5G). In contrast, FG follicles at 21% O_2_ exhibited a rapid and pronounced increase in Inhibin B (day 14: p = 0.0025; day 21: p = 0.054), closely resembling progesterone production patterns (Fig. 5D). FG follicles cultured at reduced oxygen tensions (3% and 5% O_2_) demonstrated a delayed rise in Inhibin B, consistent with observed delays in the production of progesterone and AMH (Fig. 5H).

These data strongly indicate that oxygen tension significantly modulates the timing and magnitude of steroid and peptide hormone production during follicle culture. Atmospheric oxygen conditions (21% O_2_) support rapid, early-stage increases in reproductive hormones, whereas reduced oxygen tensions (3% and 5% O_2_) induce delayed but sustained hormone production profiles. These differences likely reflect the underlying physiological adjustments of follicles to varied oxygen environments and could have critical implications for optimizing follicle growth and improving oocyte developmental outcomes in assisted reproductive technologies.

### Oxygen influences the correlation between follicle growth and hormone production

To consolidate our findings, we investigated the relationship between follicle growth and hormone production under varying oxygen tensions, aiming to better characterize follicular endocrine functionality. Regression analyses were performed on fast-growing follicles to determine the association between follicle diameter and hormone levels for E2, P4, AMH, and INHB across different oxygen tensions (Fig. 6).

**Figure 6.**
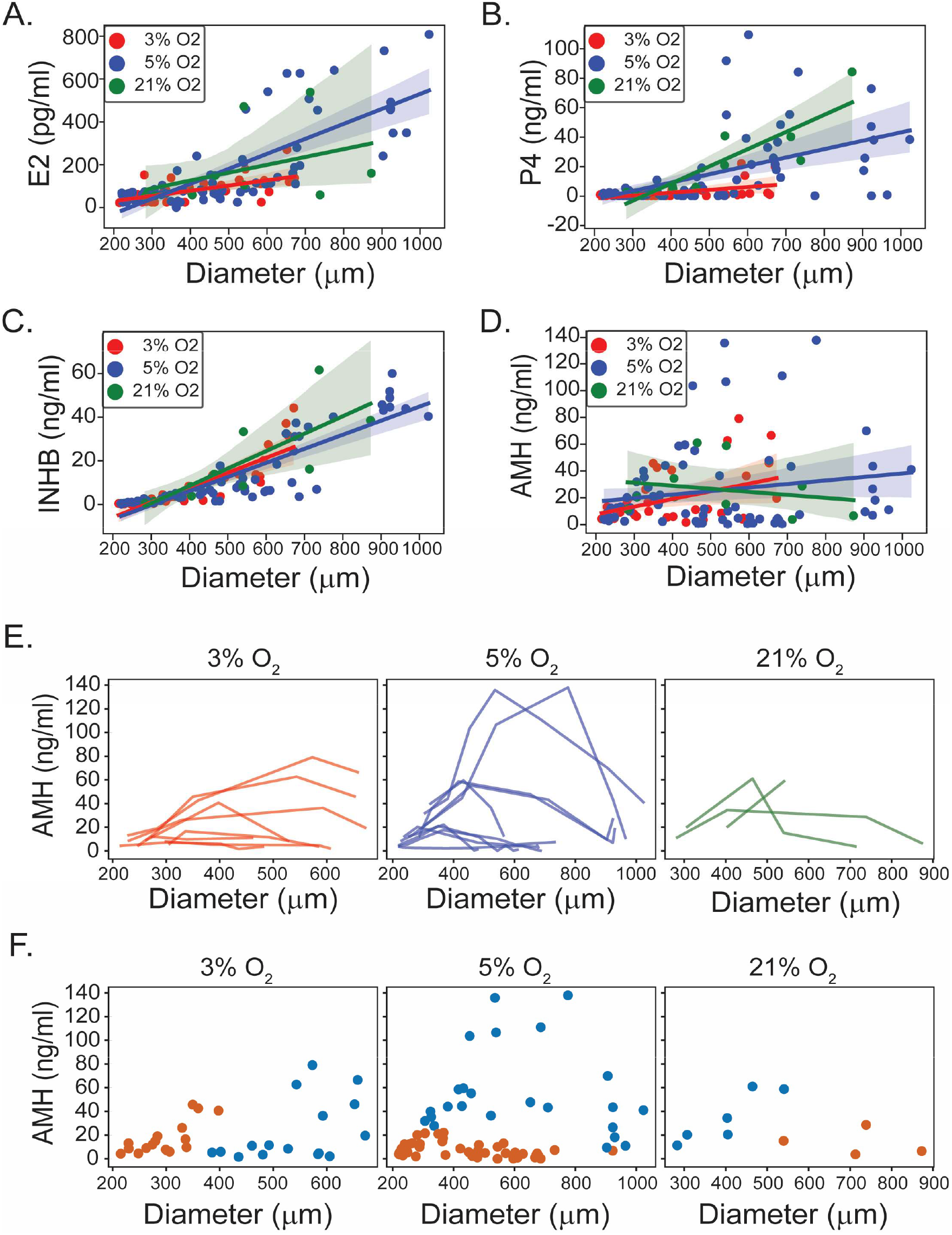
Correlations between follicle diameter and hormone production, and AMH secretion dynamics across oxygen tensions. (**A–D**) Scatter plots with regression lines illustrating correlations between follicle diameter (μm) and hormone concentrations for (A) Estradiol, (B) Progesterone, (C) Inhibin B, and (D) AMH under 3% (red), 5% (blue), and 21% O_2_ (green). Shaded areas represent 95% confidence intervals. Regression analyses revealed significant positive correlations at reduced oxygen tensions for Estradiol at 5% O_2_ (p = 0.007), Progesterone at 3% O_2_ (p = 0.003) and 5% O_2_ (p < 0.001), and Inhibin B at 3% O_2_ (p = 0.016) and 5% O_2_ (p < 0.001). No significant correlations were found under atmospheric oxygen (21% O_2_) for Estradiol (p = 0.211), Progesterone (p = 0.209), or Inhibin B (p = 0.174). AMH did not exhibit significant correlations with follicle diameter at any oxygen level (3% O_2_: p = 0.214, 5% O_2_: p = 0.339, 21% O_2_: p = 0.655). (**E**) Individual AMH secretion trajectories plotted against follicle diameter, color-coded by oxygen tension. Each line represents a single fast-growing follicle, illustrating considerable temporal heterogeneity in AMH expression independent of size. (**F**) Gaussian Mixture Modeling (GMM) clustering of the joint distribution of follicle diameter and AMH concentration under each oxygen condition, supporting the presence of discrete secretion states under all oxygen conditions, particularly at 5% O_2_.

For E2, a significant positive correlation between follicle diameter and hormone concentration was observed specifically at 5% O_2_ (p = 0.007; Fig. 6A). This indicates that larger follicles cultured under intermediate oxygen conditions (5% O_2_) were associated with increased E2 production. No significant correlations were observed at 3% (p = 0.217) or 21% O_2_ (p = 0.211). Progesterone production demonstrated robust positive correlations with follicle diameter at both 3% O_2_ (p = 0.003) and 5% O_2_ (p < 0.001; Fig. 6B). The steep slopes of regression lines at these lower oxygen tensions underscore that follicular enlargement in reduced oxygen environments was closely associated with enhanced progesterone synthesis. Conversely, this relationship was weaker and not statistically significant at 21% O_2_ (p = 0.209). Inhibin B showed significant positive correlations with follicle diameter at reduced oxygen tensions, specifically at 3% O_2_ (p = 0.016) and 5% O_2_ (p < 0.001; Fig. 6C). The regression lines for these conditions exhibited notably sharper increases compared to atmospheric conditions (21% O_2_: p = 0.174). This indicates that follicles cultured in hypoxic conditions achieve higher Inhibin B secretion in association with their growth, compared to follicles cultured in standard atmospheric oxygen.

To assess the relationship between follicle diameter and AMH secretion under different oxygen tensions, we first applied linear regression within each oxygen condition. AMH did not exhibit statistically significant correlations with follicle diameter at any oxygen level (3% O_2_: p = 0.214, 5% O_2_: p = 0.339, 21% O_2_: p = 0.655; Fig. 6D). We further explored whether polynomial regression models could better explain the data. At 3% O_2_, the linear model yielded a slightly higher adjusted R^2^ (0.130) than the quadratic model (0.105), indicating no added benefit from including a nonlinear term. At 5% and 21% O_2_, although quadratic fits marginally increased the adjusted R^2^ compared to linear models (0.0169 to 0.019, and −0.071 to 0.169, respectively), the improvements were not statistically significant (ANOVA: p = 0.290 and 0.112, respectively). In all cases, the quadratic terms were not individually significant (p > 0.1). These analyses indicate that neither linear nor polynomial models adequately capture the observed variability in AMH secretion, suggesting that AMH dynamics are not simply size-dependent but likely reflect stage-specific regulation. Interestingly, when we examined individual follicle trajectories (Fig. 6E), we observed substantial heterogeneity in AMH expression among follicles of similar diameter—some exhibiting robust secretion while others produced little to none. Rather than following a simple size-dependent trend, AMH expression appeared temporally dynamic, with secretion peaks emerging at distinct windows during follicular development. This pattern of heterogeneity was consistently evident across all oxygen conditions. To further explore potential AMH expression states, we applied Gaussian Mixture Modeling (GMM) to the joint distribution of follicle diameter and AMH levels. Model selection using the Bayesian Information Criterion (BIC) supported a three-component model at 5% O_2_ (ΔBIC = 46.6), while two-component models best fit the data for 3% and 21% O_2_ (ΔBIC = 2.4 and 0.1, respectively). These GMM-derived clusters (Fig. 6F) may reflect follicles transitioning through early, peak, or declining phases of AMH secretion, rather than representing size-based phenotypes. This interpretation is further supported by longitudinal hormone profiles, which revealed discrete and stage-specific AMH peaks during follicular growth (Fig. 5F). Together, our findings suggest that AMH serves as a temporally regulated marker of granulosa cell activity and follicular health, rather than a continuous correlate of follicle size.

Overall, our findings demonstrate a significant and oxygen-dependent linkage between follicle growth dynamics and hormone production. These data highlight oxygen tension as a critical determinant influencing follicular endocrine activity, providing key insights that may guide the optimization of culture conditions to enhance follicle developmental outcomes and hormonal function in assisted reproductive technologies.

## DISCUSSION

This study provides compelling evidence that culturing rhesus macaque secondary follicles under reduced oxygen tensions (3% and 5% O_2_) significantly improves follicle survival, growth dynamics, and hormone production compared to atmospheric oxygen (21% O_2_). Overall, follicles cultured in 3% and 5% O_2_ demonstrated similarly favorable outcomes in terms of survival, antrum formation, and growth. These findings highlight the critical importance of closely replicating physiological oxygen conditions during in vitro follicle culture to enhance follicular development and functionality.

The substantially higher survival rates observed under low oxygen conditions (3% and 5% O_2_) compared to atmospheric conditions align with previous studies demonstrating beneficial effects of reduced oxygen (5% O_2_) on follicle viability [8, 12]. Our results extend these observations, confirming the benefit of even lower oxygen tension (3% O_2_). The detrimental effects of culturing at 21% O_2_ likely arise from increased oxidative stress, known to cause cellular damage and apoptosis in granulosa cells and oocytes [9, 15]. Lower oxygen tensions presumably reduce oxidative damage, thereby creating a protective microenvironment conducive to follicle survival. Additionally, antrum formation—a critical developmental milestone—was significantly enhanced at lower oxygen tensions (3% and 5% O_2_), consistent with mammalian studies demonstrating improved antrum development and oocyte competence at reduced oxygen levels (5% O_2_) [10, 12]. Thus, lowering oxygen levels in follicle culture not only prolongs follicle survival but actively promotes key structural developmental milestones essential for successful oocyte maturation.

The distinct growth patterns we observed across different oxygen tensions yield valuable insights into the physiological regulation of follicular development. Follicles cultured at lower oxygen tensions (3% and 5% O_2_) exhibited sustained, robust growth dynamics compared to follicles at atmospheric conditions, which initially grew rapidly but subsequently degenerated. These observations reinforce the importance of mimicking *in vivo* oxygen concentrations (approximately 1.5–8.7%), which are significantly lower than standard laboratory conditions [5, 6]. Rapid growth followed by degeneration under atmospheric oxygen highlights oxidative stress as a probable contributing factor to reduced follicular longevity and functionality. Lower oxygen conditions likely enhance metabolic efficiency and follicular resilience. Intriguingly, the unique emergence of a “delayed Fast-grow” phenotype exclusively in 5% O_2_ suggests that intermediate hypoxic conditions might activate distinct developmental pathways or adaptive responses. Further mechanistic investigation into this delayed growth phenotype could yield novel insights into follicle biology.

Our results clearly illustrate that oxygen tension markedly influences steroid and peptide hormone production by follicles in vitro. Follicles cultured at 21% O_2_ showed significantly elevated early-stage estradiol and progesterone production but experienced compromised long-term viability. This initial hormone surge likely reflects increased activity of oxygen-dependent steroidogenic enzymes (e.g., aromatase [25]) or enzymes dependent on mitochondrial respiration to regenerate NAD+ (e.g. 3β-hydroxysteroid dehydrogenase [26]). The increased substrate availability at 21% O_2_ is potentially confounded by elevated oxidative stress, with impaired follicular development over time [15]. In contrast, follicles cultured at 3% and 5% O_2_ exhibited delayed but sustained elevations in estradiol and progesterone, correlating with improved survival and developmental trajectories. These findings suggest that reduced oxygen tensions maintain balanced steroidogenesis and cellular integrity, presumably by limiting oxidative stress and promoting a more physiologically relevant environment. AMH levels were relatively stable but slightly elevated under 21% O_2_ early in culture, whereas lower oxygen tensions delayed and broadened AMH production peaks, consistent with the increased expression of AMH after follicle activation that peaks in tertiary and small antral follicles [27]. Inhibin B similarly demonstrated delayed but sustained production at reduced oxygen levels. Collectively, these hormone dynamics reinforce the critical role of oxygen tension in modulating follicular endocrine function and emphasize the potential benefits of using lower oxygen tensions in reproductive biotechnology.

The present findings have significant practical implications for enhancing follicle culture systems and assisted reproductive technologies (ART). By demonstrating superior outcomes at low oxygen tensions (3% and 5% O_2_) compared to standard atmospheric conditions, our data provide strong rationale for adopting physiologically relevant, reduced-oxygen environments in clinical and research applications. Optimizing oxygen conditions could significantly improve the yield and quality of mature oocytes from follicle culture, particularly for fertility preservation strategies and *in vitro* fertilization (IVF) procedures. Such improvements are especially critical for patients with limited ovarian reserves or those undergoing ovarian tissue cryopreservation prior to gonadotoxic therapies.

While our study provides valuable insights, several limitations should be addressed in future research. Firstly, the culture duration at 3% O_2_ was limited to 7 weeks due to concerns about maintaining sterility of the cultures, and extending this period could reveal additional long–term effects. With careful maintenance of the interior environment of the hypoxia chamber, we successfully demonstrated that prolonged culture in a hypoxic glove–box incubator can be achieved without fungal or bacterial contamination. Secondly, investigating the molecular mechanisms underlying the observed differences in follicular development and hormone production across oxygen tensions would provide deeper insights into follicle physiology. The 5% and 21% O_2_ experiments were allowed to proceed to follicle failure, which prevented the molecular analysis of viable follicles. Future studies should also examine the developmental competence of oocytes matured under these conditions through *in vitro* fertilization and embryo culture experiments, with molecular and metabolic analyses at all stages of development. Additionally, exploring even lower oxygen tensions (1–2% O_2_), may more closely mimic the oxygen levels found in the relatively avascular cortex [28, 29]. Integrating dynamic oxygen gradients to increase oxygen concentrations as the follicles progress from the secondary to the antral stages may better replicate the changing oxygen environment experienced by follicles as the ovarian vasculature expands to feed the growing antral follicle pool [30].

In conclusion, our study demonstrates that lower oxygen tensions (3% and 5% O_2_) markedly improve rhesus macaque follicle survival, growth dynamics, and endocrine function compared to atmospheric oxygen (21% O_2_). These findings highlight the essential role of physiologically relevant oxygen conditions for optimizing follicle culture systems. Through improved replication of the ovarian microenvironment, reduced oxygen culture conditions offer a promising strategy to enhance developmental outcomes in ART. Ultimately, this approach could significantly improve fertility preservation methods and clinical IVF outcomes, benefiting patients with diminished ovarian reserves or those at risk of infertility due to medical treatments.

## Abbreviations

IVFD: *in vitro* follicle development
E2: estradiol
P4: progesterone
AMH: anti–Müllerian hormone
ART: assisted reproductive technologies;

## Acknowledgments

We are grateful to David Erickson and the staff of the ONPRC Endocrine Technologies Core for conducting hormone measurements. The ETC is partially supported by NIH grant P51 OD011092 for operation of the Oregon National Primate Research Center. Research reported in this publication was supported by the National Institutes of Health Center for Children’s Health and Disease (R01 HD082208 and R01 HD083930 to AJK and MZ, respectively). The content is solely the responsibility of the authors and does not necessarily represent the official views of the NIH.

## Author Contributions

KW, SW, MBZ, and AJK contributed to study design, conducted experiments, interpreted data, and contributed to manuscript writing and editing.

All data generated or analyzed during this study are included in this article.

